# Human gut commensal *Alistipes timonensis* modulates the host lipidome and delivers anti-inflammatory outer membrane vesicles to suppress colitis in an *Il10*-deficient mouse model

**DOI:** 10.1101/2024.10.23.619966

**Authors:** Ethan A. Older, Mary K. Mitchell, Andrew Campbell, Xiaoying Lian, Michael Madden, Yuzhen Wang, Lauren E. van de Wal, Thelma Zaw, Brandon N. VanderVeen, Rodney Tatum, E. Angela Murphy, Yan-Hua Chen, Daping Fan, Melissa Ellermann, Jie Li

## Abstract

Correlative studies have linked human gut microbes to specific health conditions. *Alistipes* is one such microbial genus negatively linked to inflammatory bowel disease (IBD). However, the protective role of *Alistipes* in IBD is understudied, and the underlying molecular mechanisms remain unknown. In this study, colonization of *Il10*-deficient mice with *Alistipes timonensis* DSM 27924 delays colitis development. Colonization does not significantly alter the gut microbiome composition, but instead shifts the host plasma lipidome, increasing phosphatidic acids while decreasing triglycerides. Outer membrane vesicles (OMVs) derived from *Alistipes* are detected in the plasma of colonized mice, carrying potentially immunomodulatory metabolites into the host circulatory system. Fractions of *A. timonensis* OMVs suppress LPS-induced *Il6*, *Il1b*, and *Tnfa* expression *in vitro* in murine macrophages. We detect putative bioactive lipids in the OMVs, including immunomodulatory sulfonolipids (SoLs) in the active fraction, which are also increased in the blood of colonized mice. Treating *Il10*-deficient mice with purified SoL B, a representative SoL, suppresses colitis development, suggesting its contribution to the anti-inflammatory phenotype observed with *A. timonensis* colonization. Thus, *A. timonensis* OMVs represent a potential mechanism for *Alistipes*-mediated delay of colitis in *Il10*-deficient mice via delivery of immunomodulatory lipids and modulation of the host plasma lipidome.

## Introduction

The trillions of microbes inhabiting the human gut are inherently linked to human health through interactions with their host environment. These complex interactions affect human health in various negative and positive ways; for example, *Helicobacter pylori* can lead to the formation of gastric ulcers and later gastric cancer whereas, *Akkermansia muciniphila* is inversely associated with obesity, diabetes, cardiometabolic diseases and inflammation.^1,2^ The composition of the gut microbiome is delicately balanced and it dynamically responds to external stimuli. When this balance is significantly disturbed, it can impair crucial functions such as maintaining immune homeostasis and can contribute to the development and exacerbation of various diseases including cancer, cardiovascular disease, and chronic inflammation.^3–5^ This disturbance in microbiome composition and resulting functions is known as microbial dysbiosis. Inflammatory bowel disease (IBD) is particularly affected by microbial dysbiosis.^6^ While numerous studies have reported major shifts in the gut microbiome composition with IBD pathogenesis, there is a lack of mechanistic understanding regarding how specific bacterial populations might protect against IBD progression.

Another hallmark of IBD is dysregulation of the circulating plasma lipid profile, known as dyslipidemia.^7–9^ Normally, circulating lipids in the plasma play a role in maintaining homeostasis in processes throughout the body including energy metabolism, cardiovascular health, and managing inflammation.^10,11^ Imbalances in these lipids lead to perturbations in their respective processes and often have far reaching effects.^11^ For example, dysregulation of specific lipids such as sphingosine-1-phosphate and triglycerides have been correlated with IBD pathogenesis.^12,13^ Gut microbes have also been found to modulate the host lipidomic profile and impact host physiology, further complicating the interplay between the gut microbiome and human health.^8,14^ *Alistipes* is an emerging microbial genus that is consistently found in the human gut microbiome.^15^ As a non-pathogenic commensal, *Alistipes* has shown increasing human health relevance and has been linked to protective effects against numerous inflammatory diseases by correlative gut microbial sequencing studies.^16–19^ Importantly, *Alistipes* abundance has been negatively associated with disease severity in IBD patients and its metabolites were shown to be decreased in an *Il10*-deficient mouse model of experimental colitis, supporting that *Alistipes* may play a protective role against IBD through its metabolite production.^20^ *Alistipes* colonization in the DSS and oxazolone-induced mouse models of experimental colitis has been found to attenuate inflammation through the induction of endogenous anti-inflammatory cytokines including IL-10.^21,22^ While this interaction has been documented, it remains unclear how *Alistipes* induces this phenotype and to what extent *Alistipes*-derived microbial functional metabolites mediate interactions between *Alistipes* and the host.

It has been proposed that gut bacteria secrete outer membrane vesicles (OMVs) to interact with their hosts. OMVs are spherical nanoparticles derived from the outer membranes of Gram-negative bacteria.^23^ Their size ranges between 20 – 300 nm in diameter and they are typically composed of a lipid bilayer containing phospholipids, lipopolysaccharide (LPS), and outer membrane proteins. They are deliberately produced in response to stress signals including temperature, oxidation, and nutrient availability, and are particularly important in mediating interactions among microbes, hosts, and other organisms.^24,25^ Notably, OMVs can be selectively packaged by their producers and serve as delivery vehicles for their cargo which can include metabolites, proteins, DNA, and RNA.^26–29^ This provides a means to carry microbial components through the protective mucus layer and enable otherwise gut-restricted elements to cross the colonic epithelium and reach host cells via the cardiovascular system, initiating various host responses.^28,30–32^

Here, we show that colonization of *Il10*-deficient mice with *Alistipes timonensis* DSM 27924, a producer of sulfonolipids (SoLs), delays the development of colitis. We show that *Alistipes* colonization does not significantly affect the gut microbiome composition during colitis development, but results in a shift of the host plasma lipidome consistent with clinical observations. We find OMVs derived from *A. timonensis* in the blood plasma of colonized mice which carry potentially immunomodulatory *A. timonensis* metabolites to the host circulatory system. We then show that fractionated OMVs stimulate an anti-inflammatory response *in vitro* in murine macrophages and we detect immunomodulatory SoLs in the active fraction. We finally demonstrate that a purified SoL B fraction isolated from *A. timonensis* OMVs also delays colitis in *Il10*-deficient mice. Taken together, this study reports for the first time that *A. timonensis* delays colitis in the *Il10*-deficient mouse model and proposes *A. timonensis*-derived SoL B, which is delivered to host circulation by OMVs, as a potentially contributing factor to the anti-inflammatory action of *A. timonensis* colonization.

## Results

### Colonization of *Alistipes timonensis* DSM 27924 in *Il10*-deficient mice delayed colitis progression

To determine whether *Alistipes* impacts intestinal inflammation development, we utilized the well-established *Il10*-deficient (IL10 KO) mouse model of piroxicam-accelerated colitis.^33–35^ This mouse model captures the genetic and environmental interactions and contributions that drive IBD development.^34^ Further, *Alistipes* colonization in alternative models of experimental colitis has shown that colonization induces innate IL-10 production which is a major contributor to anti-inflammatory phenotypes.^21,22^ Thus, the use of the IL10 KO model will allow us to examine the effects of *Alistipes* and its metabolites on colitis progression without influence from endogenous IL-10 production. *Alistipes timonensis* DSM 27924 was engrafted into the gut microbiomes of the colitis-susceptible IL10 KO mice at the start of colitis induction via a diet supplemented with piroxicam. *A. timonensis* DSM 27924 was selected because it was previously shown to colonize the gut of mice.^36^ Additionally, while the genus *Alistipes* has been negatively correlated with inflammatory diseases, such as IBD, this species of *Alistipes* has not been previously positively or negatively linked to any disease.^16,20^

Two groups of female IL10 KO mice were used: group 1, piroxicam (uncolonized) and group 2, piroxicam + *A. timonensis* (*A. timonensis*-colonized). A confirmatory second independent cohort of female IL10 KO mice was also used, and all results were found to be consistent between the two cohorts. Thus, the data from these two independent cohorts were pooled for all further analysis. PCR amplification of the *A. timonensis* 16S rRNA gene confirmed the presence and persistence of *A. timonensis* after oral gavage, supporting successful colonization and confirming that *A. timonensis* was not present prior to engraftment (**Supplemental Figure S1a**). Uncolonized mice developed colitis as indicated by decreased body weight over time (**Supplemental Figure S1b,c**) and increased gross pathology and histology scores at necropsy (**Figure 1a**). In contrast, *A. timonensis*-colonized mice showed reduced weight loss compared to uncolonized mice as well as significantly decreased gross pathology and histology scores (**Supplemental Figure S1b,c**) (**Figure 1a**), suggesting that *A. timonensis* may act to suppress colitis development in the IL10 KO mice. Consistent with model expectations, gross pathology and histology scoring indicated that suppression of intestinal inflammation was localized to the colon and was more pronounced in the distal colon (**Supplemental Files 1 and 2**).^33,34^ Inflammatory cytokine profiling of RNA extracted from the distal colon via reverse transcription quantitative PCR (RT-qPCR) revealed that *A. timonensis*-colonized mice had decreased expression of pro-inflammatory cytokines *Il1b*, *Tnfa*, and *Il6*, as well as pro-inflammatory marker *Nos2* (**Figure 1b-e**), further supporting that *A. timonensis* colonization suppressed intestinal inflammation.

**Figure 1.**
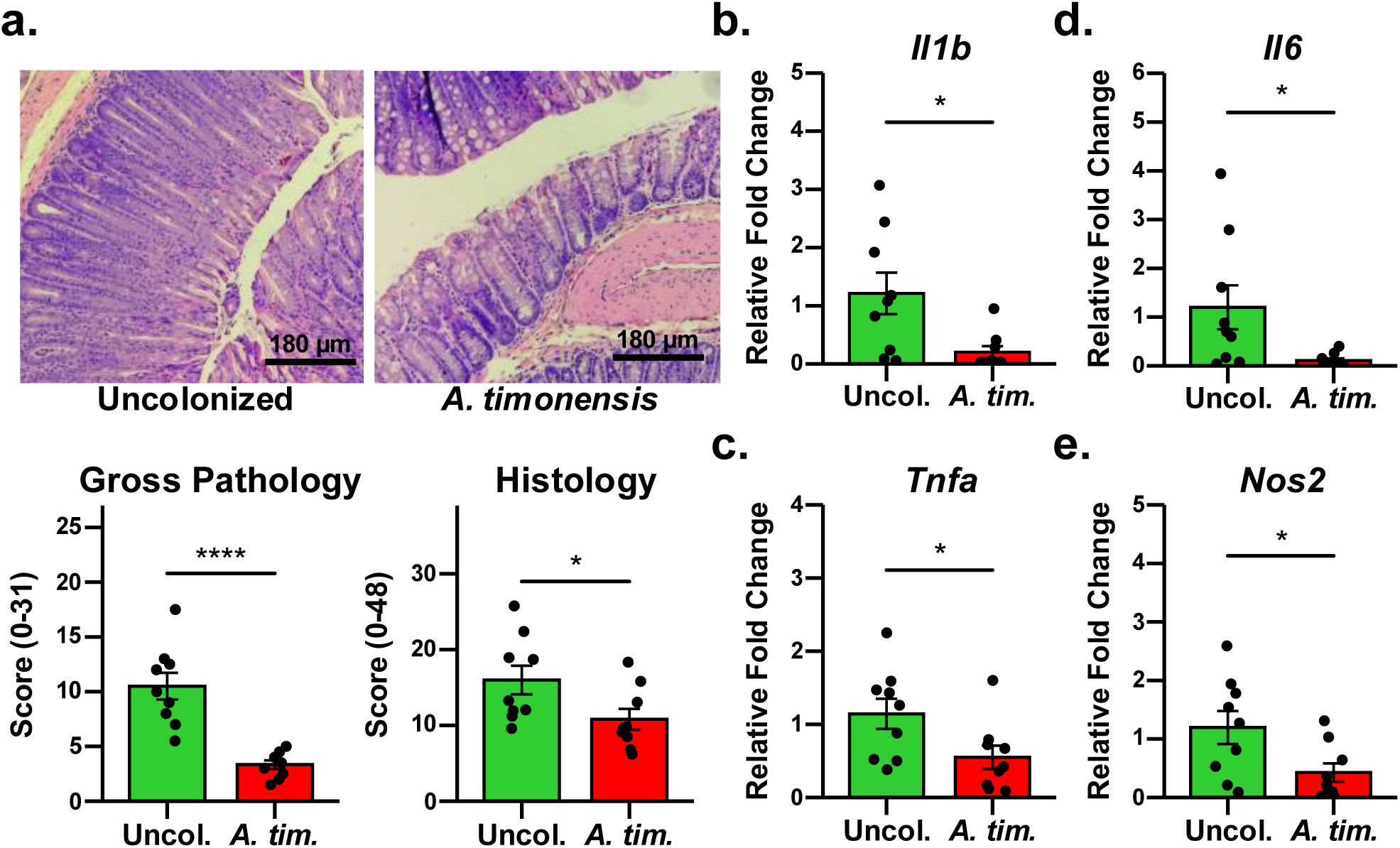
*A. timonensis* colonization delays piroxicam-accelerated colitis development in IL10 KO mice. (a) Representative H&E images of uncolonized (Uncol.) and *A. timonensis*-colonized (*A. tim.*) mouse colons showing less inflammation in colonized mice. One-sided Student’s *t*-test confirmed significantly decreased gross pathology and histology scores in Uncol. (*n* = 9 mice) compared to *A. tim.* (*n* = 9 mice) mice. (b-e) The expression of pro-inflammatory factors *Il1b*, *Tnfa*, *Il6*, *and Nos2* were significantly decreased in the distal colons of *A. timonensis*-colonized (*A. tim., n* = 9 mice) mice compared to uncolonized (Uncol.*, n* = 9 mice) mice, supporting that *Alistipes* colonization mediated a suppressive effect against colitis progression. Significance was determined using two-sided Student’s *t*-test. For all *p*-values, * 0.05 > *p* > 0.0 and **** 0.0001 > *p*.

Colitis in this model is driven by enhanced colonic epithelial apoptosis leading to a loss of gut barrier integrity.^34^ Thus, we examined cellular tight junctions using immunofluorescence microscopy and found that tight junction morphology was restored in *A. timonensis*-colonized colon samples (**Figure 2a**). We also measured the expression of tight junction proteins, revealing that *A. timonensis*-colonized mice had significantly increased expression of *claudin-3* concurrent with decreased expression of *claudin-7*. (**Figure 2b**). Claudin-3 has been previously reported to decrease in colitis, reflecting a loss of gut barrier integrity and leakage of gut contents into the host circulation.^37^ Claudin-7 has been linked to intestinal epithelial stem cell renewal and regulation and its decreased expression in *A. timonensis*-colonized mice here may suggest a reduced need for cellular repair in the epithelium due to reduced colitis after *A. timonensis* colonization.^38^

**Figure 2.**
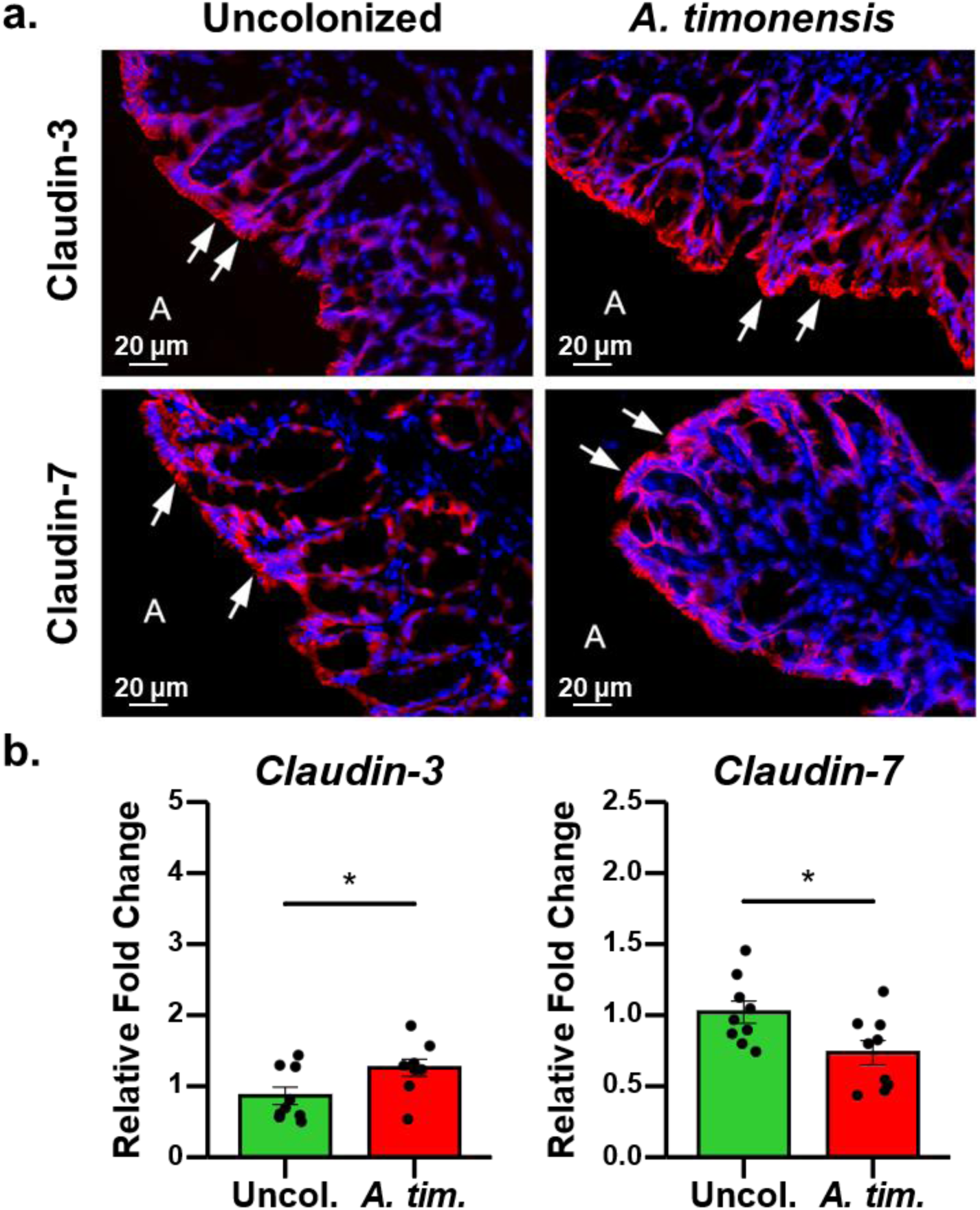
*A. timonensis* colonization modulates tight junction protein morphology and gene expression in IL10 KO mouse colons. (a) Immunofluorescent staining of tight junction proteins claudin-3 and claudin-7 (red) illustrating restoration of tight junction morphology and reduction of tight junction repair with *A. timonensis* colonization, respectively. Nuclei were stained with DAPI (blue). A: Apical side of colon tissue; Arrow: tight junction morphologies. (b) Gene expression of *claudin-3* and *claudin-7* in uncolonized (Uncol., *n* = 9 mice) and *A. timonensis*-colonized (*A. tim.*, *n* = 9 mice) mice, which is consistent with immunofluorescence staining. Significance was determined using two-sided Student’s *t*-test. For all *p*-values, * 0.05 > *p* > 0.01.

### The anti-inflammatory effects of *A. timonensis* in IL10 KO mice are independent from reversal of microbial dysbiosis

Gut dysbiosis is a hallmark of IBD, with prolonged dysbiosis leading to reduced intestinal homeostasis and ultimately chronic inflammation.^6,39^ In the IL10 KO mouse model, dysbiosis further exacerbates inflammation due to increased leakage of gut contents through the damaged colonic epithelium.^34^ Considering this, we first hypothesized that *A. timonensis* colonization may mediate an anti-colitic effect by preventing or reversing perturbations in gut microbiota composition, resulting in protection against IBD-related gut dysbiosis. Thus, we performed 16S rRNA sequencing to profile the gut microbiome composition in *A. timonensis*-colonized (*A. timonensis, n* = 10 mice) and uncolonized (Uncolonized, *n* = 11 mice) IL10 KO mice - both groups of which were treated with piroxicam. An age-matched control group of IL10 KO mice (Control, *n* = 17 mice) that were neither treated with piroxicam nor colonized with *A. timonensis* was also included (**Supplemental File 3**). Comparing uncolonized mice to control mice, microbiome composition analysis revealed a clear shift in the microbial profile (Figure 3a) accompanied by a significant decrease in alpha diversity (Figure 3b). This loss in microbial diversity is indicative of microbial dysbiosis in colitis development, consistent with the literature.^40,41^

**Figure 3.**
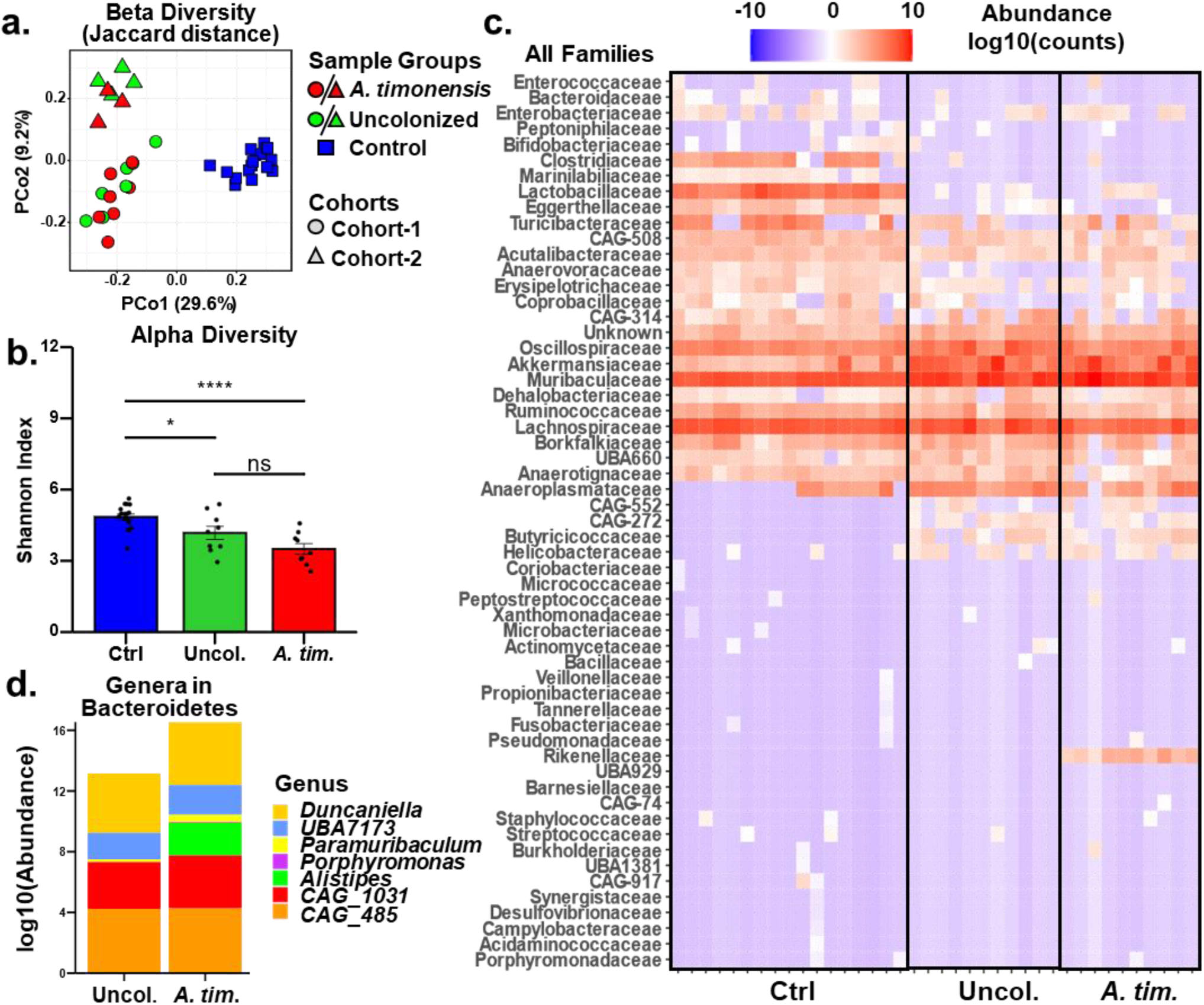
Microbiome composition analysis indicates *A. timonensis* colonization does not reverse dysbiosis. (a) Beta diversity measured by principal coordinate analysis (PCoA) using Jaccard distances shows clear separation between both independent cohorts (red and green triangles and circles) and the control group (blue squares) but no separation between *A. timonensis*-colonized (red) versus uncolonized (green) microbiome profiles. (b) Alpha diversity measured by Shannon diversity index indicates a significant loss of microbial diversity from control (*n* = 17 mice) to uncolonized (*n* = 11 mice) and *A.timonensis*-colonized (*n* = 10 mice) mice that was not restored by *A. timonensis* colonization. Significance was determined using two-sided Student’s *t*-test. For all *p*-values, * 0.05 > *p* > 0.01, ** 0.01 > *p* > 0.001, and ns: not significant. (c) Heatmap of microbiome composition at the family level showing a clear difference in control samples compared to uncolonized and *A. timonensis*-colonized mice. (d) Genera composition of the phylum Bacteroidetes highlights the major expansion of *A. timonensis* in colonized compared to uncolonized samples.

However, despite the attenuated inflammation observed with *A. timonensis* engraftment, the microbiota profile of *A. timonensis*-colonized mice showed no clear distinction from that of uncolonized mice and had no significant change in alpha diversity compared to uncolonized mice (**Figure 3a,b**). To verify this observation, we analyzed the second cohort independently and observed a consistent lack of significant microbiome compositional changes between *A. timonensis*-colonized and uncolonized mice (**Figure 3a**). At the family level, we observed clear differences between Control samples and both uncolonized and *A. timonensis*-colonized samples but observed only minor changes between uncolonized and *A. timonensis*-colonized samples, with one prominent family that *A. timonensis* belongs to, Rikenellaceae, notably more abundant in *A. timonensis*-colonized samples (**Figure 3c**). Since *Alistipes* is a member of the phylum Bacteroidetes, we also looked at changes in genera within Bacteroidetes; we observed that an increase in the abundance of *Alistipes* in *A. timonensis*-colonized compared to uncolonized samples was the only major difference (**Figure 3d**). This confirmed that the only major change in the gut microbiome profile upon *A. timonensis* colonization was an expected increase in *Alistipes*.

In contrast with many other gut microbes such as *Bacteroides uniformis* that has been shown to reduce colitis through significant modulation of the gut microbiome composition when colonized, colonization with *A. timonensis* appears to mediate its anti-colitic effect independently of major microbiome remodeling.^42^ This unexpected yet intriguing result suggests that *A. timonensis* itself plays a more direct role in the observed anti-colitic effect and focused the study towards exploring more direct host-bacterial mechanisms by which this strain modulates host inflammation.

### *A. timonensis* colonization modulates host lipidomic profile associated with colitis

Considering that the observed anti-colitic phenotype was reflected in host cytokine and host tight junction protein and gene expression, we hypothesized that *A. timonensis* colonization may directly induce a change in host signaling pathways. Notably, host immune signaling pathways are tightly regulated by circulating blood immunomodulatory lipids.^8,43^ The levels of these lipids are directly affected by the gut microbiome in various ways, such as through direct microbial biotransformation of primary bile acids to secondary bile acids or through bacterial sphingolipid-dependent downregulation of endogenous sphingolipid biosynthesis as well as the production and circulation of other microbial metabolites.^44,14^ To examine the effects of *A. timonensis* colonization on circulating host lipids, lipidomic analysis of the mouse blood plasma was conducted (**Supplemental File 4**). Using orthogonal partial least squares discriminant analysis, we observed that the host lipid profile was significantly different between *A. timonensis*-colonized and uncolonized mice (**Figure 4a**). Among the significantly changed lipids, phospholipids were found to be more abundant in *A. timonensis*-colonized compared to uncolonized samples, while triglycerides were more abundant in uncolonized compared to *A. timonensis*-colonized samples. (**Figure 4b**). These results were consistent with clinical observations that triglycerides are increased in both ulcerative colitis and Crohn’s disease.^13^

**Figure 4.**
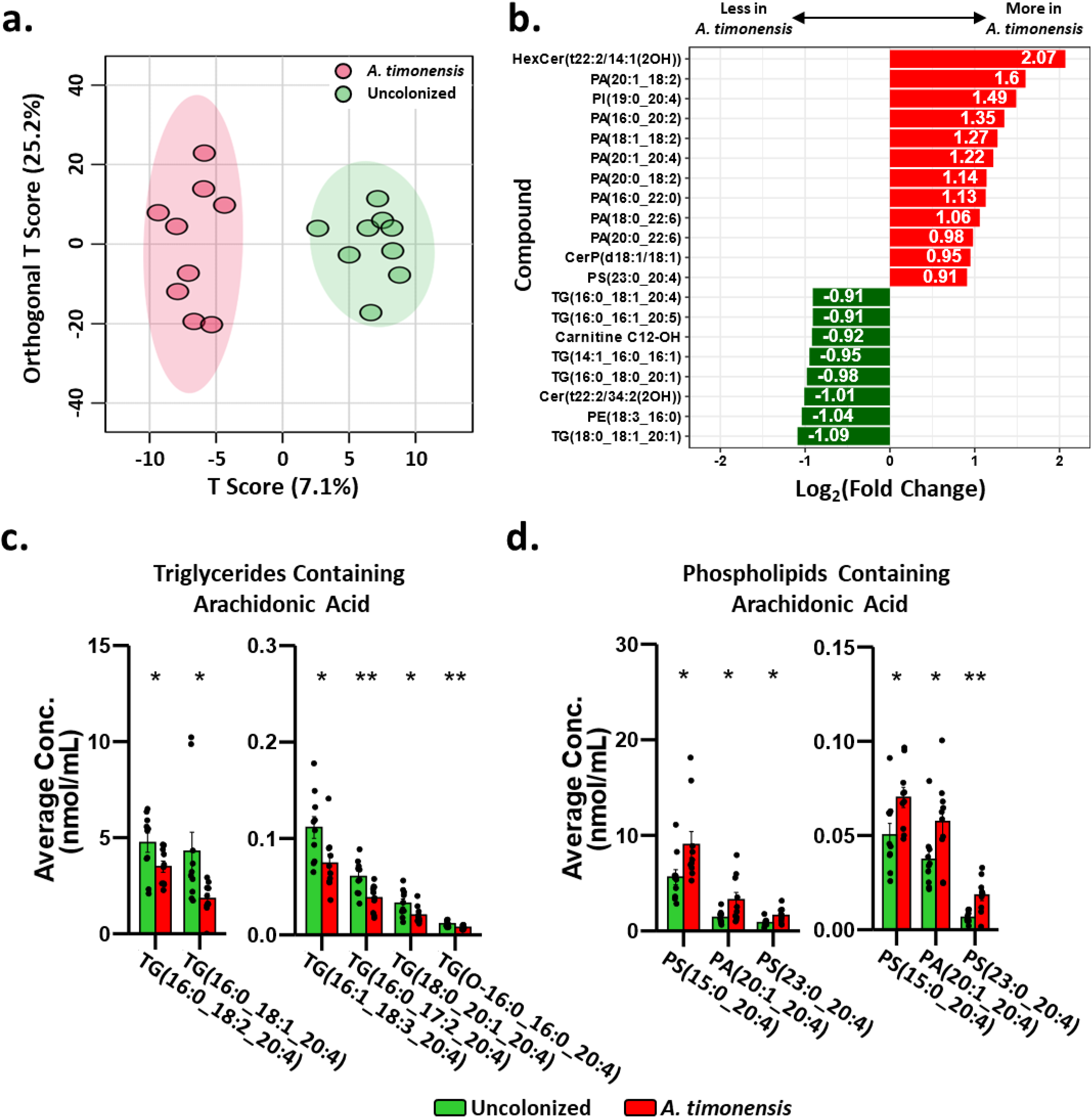
Plasma lipidomics reveals colonization-driven change in circulating lipid profile. (a) Orthogonal partial least squares-discriminant analysis (OPLS-DA) shows clear difference in plasma lipid profiles between uncolonized (*n* = 9 mice) and *A. timonensis*-colonized (*n* = 9 mice) mice. (b) The top 20 significantly differentially abundant lipids reveal a marked depletion of triglycerides (TGs) and accumulation of phosphatidic acids (PAs) in *A. timonensis*-colonized mice compared to uncolonized mice. (c,d) *A. timonensis* colonization affected the balance of two pools of arachidonic acid-containing lipids: triglycerides (c), which were less abundant in *A. timonensis*-colonized mice, and phospholipids (d), which were more abundant in *A. timonensis*-colonized mice. Significance was determined using two-sided Student’s *t*-test. For all *p*-values, * 0.05 > *p* > 0.01 and ** 0.01 > *p* > 0.001.

A recent study in acutely stimulated immune cells demonstrated that arachidonic acid (AA) is mobilized first from membrane phospholipids, which is then replenished by triacylglycerol AA after inflammation has subsided.^45^ We observed a similar trend in that AA-containing phosphatidic acids were accumulated in *A. timonensis*-colonized compared to uncolonized samples while triglycerides were decreased in *A. timonensis*-colonized samples (**Figure 4c,d**) (**Supplemental Table 1**). This may represent the replenishment of AA from triglycerides to phospholipids in association with an *A. timonensis*-mediated suppression of the inflammatory response following piroxicam challenge. These results indicate that *A. timonensis* colonization indeed modulates host signaling pathways by affecting immunomodulatory lipid regulation and suggests that a factor derived from *A. timonensis* may be involved in mediating this effect. To explore this potential factor from a molecular point of view, we considered the production and translocation of metabolites by *A. timonensis*.

### *A. timonensis* secreted outer membrane vesicles (OMVs) bearing biologically active lipids into the plasma after colonization

OMVs are ubiquitously produced by Gram-negative bacteria and are known to carry bacterially-derived elements away from the producing cell.^27,28,31^ These bacterial delivery vehicles represent a potential mechanism for metabolite translocation to the host circulation. Thus, we collected the plasma of uncolonized and *A. timonensis*-colonized mice and purified OMVs using density gradient ultracentrifugation (**Supplemental Figure S2**). To ensure detection, aliquots of plasma from each mouse were pooled together into representative samples. Indeed, OMV-like nanoparticles were purified from the pooled plasma and visualized by transmission electron microscopy (TEM) (**Figure 5a**) (**Supplemental Figure S3**), suggesting the presence of OMVs in the samples. To determine if these OMVs were related to *A. timonensis* colonization, we used dynamic light scattering (DLS) to measure the size distributions of OMVs isolated from the plasma of uncolonized and *A. timonensis*-colonized mice in comparison to OMVs isolated from a monoculture of *A. timonensis*. DLS showed that OMVs from plasma are distributed into two distinct populations (10-55 nm and 55-330 nm) and that OMVs from *A. timonensis*-colonzied mouse plasma showed an increased abundance of particles corresponding to the diameter range matching that of OMVs from *A. timonensis* culture (**Figure 5b**). This data indicates that *A. timonensis* colonization resulted in more OMVs in the plasma which are potentially derived from *A. timonensis.* To confirm that these OMVs came from *A. timonensis*, we conducted targeted metabolomic analysis looking for biomarkers of *A. timonensis*. We and others have previously shown that sulfonolipids (SoLs) are a class of molecules that are uniquely produced by the gut microbes *Alistipes* and *Odoribacter*.^35,36,46^ SoLs are further known to localize to the outer membrane of producing bacteria,^47^ strongly suggesting they may be found in OMVs. Indeed, our metabolomic analysis revealed a significantly higher abundance of sulfobacin B (SoL B), the major SoL produced by *Alistipes*,^35^ in *A. timonensis*-colonized plasma OMV samples compared to uncolonized plasma OMV samples (**Figure 5c**). Considering our DLS results, this metabolomic evidence, and our microbiota composition analysis which showed that increased abundance of SoL-producing *A. timonensis* was the only major change after colonization, the OMVs we isolated from the plasma of *A. timonensis*-colonized mice likely contain an abundance of *A. timonensis*-derived OMVs.

**Figure 5.**
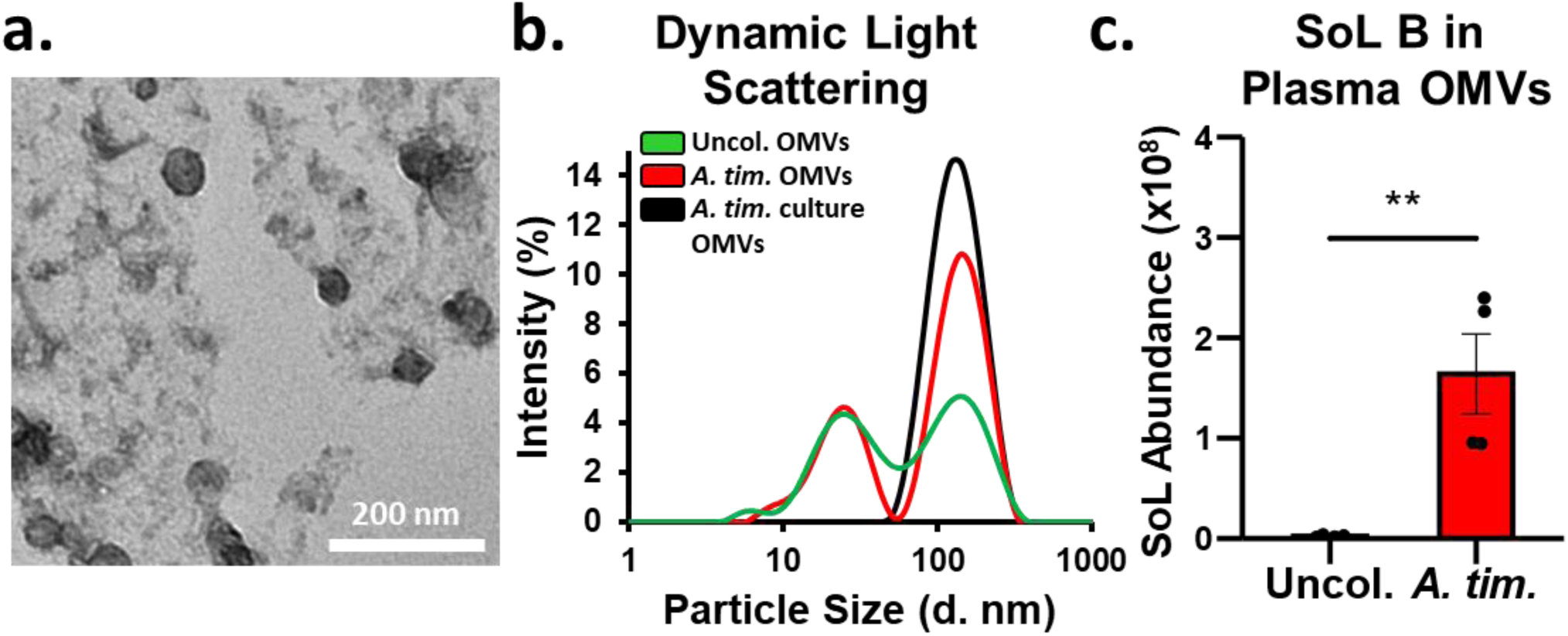
OMVs purified from the plasma of *A. timonensis*-colonized mice are abundant in *A. timonensis*-derived OMVs. (a) TEM was used to visually confirm the presence of OMVs after negative staining. x30.0k magnification was used to confirm the diameter of OMVs in the expected range. (b) DLS was used to measure the size distribution of plasma OMVs from uncolonized (green) and *A. timonensis*-colonized (red) mice alongside OMVs purified from *A. timonensis* culture (black). DLS analysis showed two clear vesicle populations with diameter ranges of 10-55 nm and 55-320 nm. The rightmost larger diameter population corresponds with *A. timonensis* culture OMVs and reflects the increased abundance of OMVs derived from *A. timonensis* colonization. (c) Lipids were extracted from purified blood OMVs, measured using targeted metabolomics, and SoL B was found to be significantly increased in plasma OMVs from *A. timonensis*-colonized (*n* = 4 mice) compared to uncolonized (*n* = 4 mice) mice. Significance was determined using two-sided Student’s *t*-test. For all *p*-values, ** 0.01 > *p* > 0.001.

Microbial metabolites tend to be restricted within the gut if they are not actively delivered to host circulation.^31,48^ Since the OMVs were found to carry SoL B, we considered if *A. timonensis*-derived lipids, including SoLB, could be delivered to the blood of mice after *A. timonensis* colonization. Accordingly, targeted metabolomic analysis of the host plasma lipidome showed a significant increase of SoL B in the plasma and also indicated a significant increase in the abundance of sulfobacins A (SoL A) and F (SoL F) in *A. timonensis*-colonized compared to uncolonized plasma (**Figure 6a-d**). The identity of SoLs was confirmed in these samples by retention time and MS/MS fragmentation matching compared to in-house standards and literature references (**Figure 6e,f**).^49^ This result suggested that *A. timonensis*-derived OMVs indeed delivered *A. timonensis*-derived metabolites including SoLs to the host circulatory system. The delivery of these non-gut restricted lipids thus represents a potential method by which *A. timonensis* colonization suppresses the colitis phenotype. To support this, we considered if the increased abundance of SoLs in the plasma was due to passive diffusion or if *A. timonensis* OMVs containing SoLs were actively translocated to the host circulation. Since the SoL-producer *Alistipes* colonizes the gut of the mice, we assumed that the production of SoL B is primarily in the gut and thus the concentration of SoL B should be greatest in the feces. If SoL B passively diffuses into host circulation, then we would expect a higher abundance of SoL B in the feces compared to the plasma. If SoL B is actively translocated to host circulation, then we would expect the abundance of SoL B to be greater in the plasma than the feces. We compared the fold changes in SoL B abundance between fecal material and plasma in both uncolonized and *A. timonensis*-colonized samples. We found that SoL B increased by 1.8-fold in uncolonized mice while it increased by 3.9-fold in *A. timonensis*-colonzied mice (**Figure 6g**). In both cases, SoL B was greater in the plasma than in the feces, suggesting that SoL B is actively translocated from the gut into host circulation. Taken together with the presence of SoL B in *A. timonensis* OMVs and increased abundance of these OMVs in *A. timonensis*-colonized mice, it is likely that OMVs serve as a vehicle for active delivery of *A. timonensis*-derived SoL B to host circulation.

**Figure 6.**
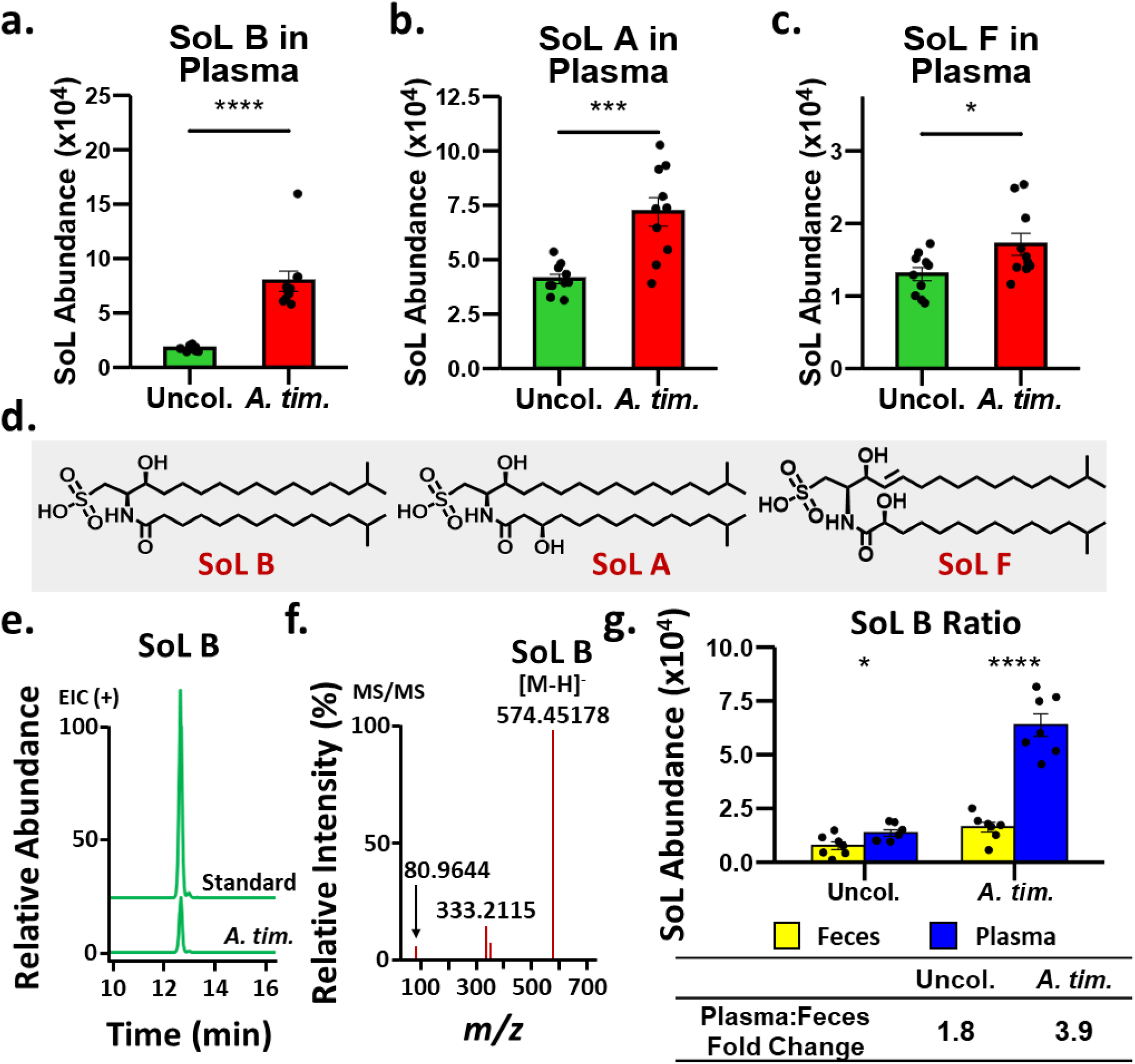
Sulfonolipid abundance significantly increased in the plasma of *A. timonensis*-colonized mice. (a-c) Abundance of SoL B (a), SoL A (b), and SoL F (c) measured in uncolonized (*n* = 10 mice) compared to *A. timonensis*-colonized (*n* = 10 mice) mouse plasma by targeted metabolomics. (d) Corresponding chemical structures of SoL B, A, and F. (e) Retention time comparison of in-house SoL B standard and observed SoL B peak in *A. tim.* samples supporting identification as SoL B. (f) MS/MS fragmentation used to positively identify SoL B based on characteristic fragmentation including the ∼81 m/z fragment corresponding to the loss of the HSO ^-^ headgroup. (g) The plasma-to-feces ratio of SoL B abundance was significantly greater in *A. timonensis*-colonized (*n* = 7) compared to uncolonized (*n* = 7) samples, suggesting that the increased production of SoL B by *A. timonensis* in the gut led to increased SoL B abundance in the plasma via active translocation of SoL B from the gut to the plasma by OMVs. Significance was determined using two-sided Student’s *t*-test. For all *p*-values, * 0.05 > *p* > 0.01, ** 0.01 > *p* > 0.001, *** 0.001 > *p* > 0.0001, and **** 0.0001 > *p*.

### *A. timonensis* OMV fraction containing bioactive lipids exerts an anti-inflammatory effect *in vitro* in murine macrophages

Because *Alistipes*-derived OMVs were found in the host circulation, we considered if these OMVs contributed to the observed anti-colitic phenotype. To obtain enough OMVs for an *in vitro* assay, we purified OMVs from a culture of *A. timonensis*. Due to the complex chemical composition of OMVs, we considered if one or more of their components possessed anti-inflammatory activity. We thus fractionated the OMV samples based on polarity and tested each fraction’s ability to suppress LPS-induced inflammation in macrophages. We observed that one fraction, Fraction 2 (F2), significantly suppressed LPS-induced inflammation as measured by fold change of expression of the cytokines *Il1b* and *Il6*, while *Tnfa* showed a similar trend (**Figure 7a**). Importantly, the leakage of gut antigens, such as LPS, into the blood, leads to immune system activation, which is one of the driving factors of colitis in the IL10 KO mouse model.^34^ Thus, the suppression of LPS-induced pro-inflammatory cytokine expression by F2 suggested that one or more components in *A. timonensis* OMVs may modulate the immune system by partially blocking the LPS-mediated inflammatory response.

**Figure 7.**
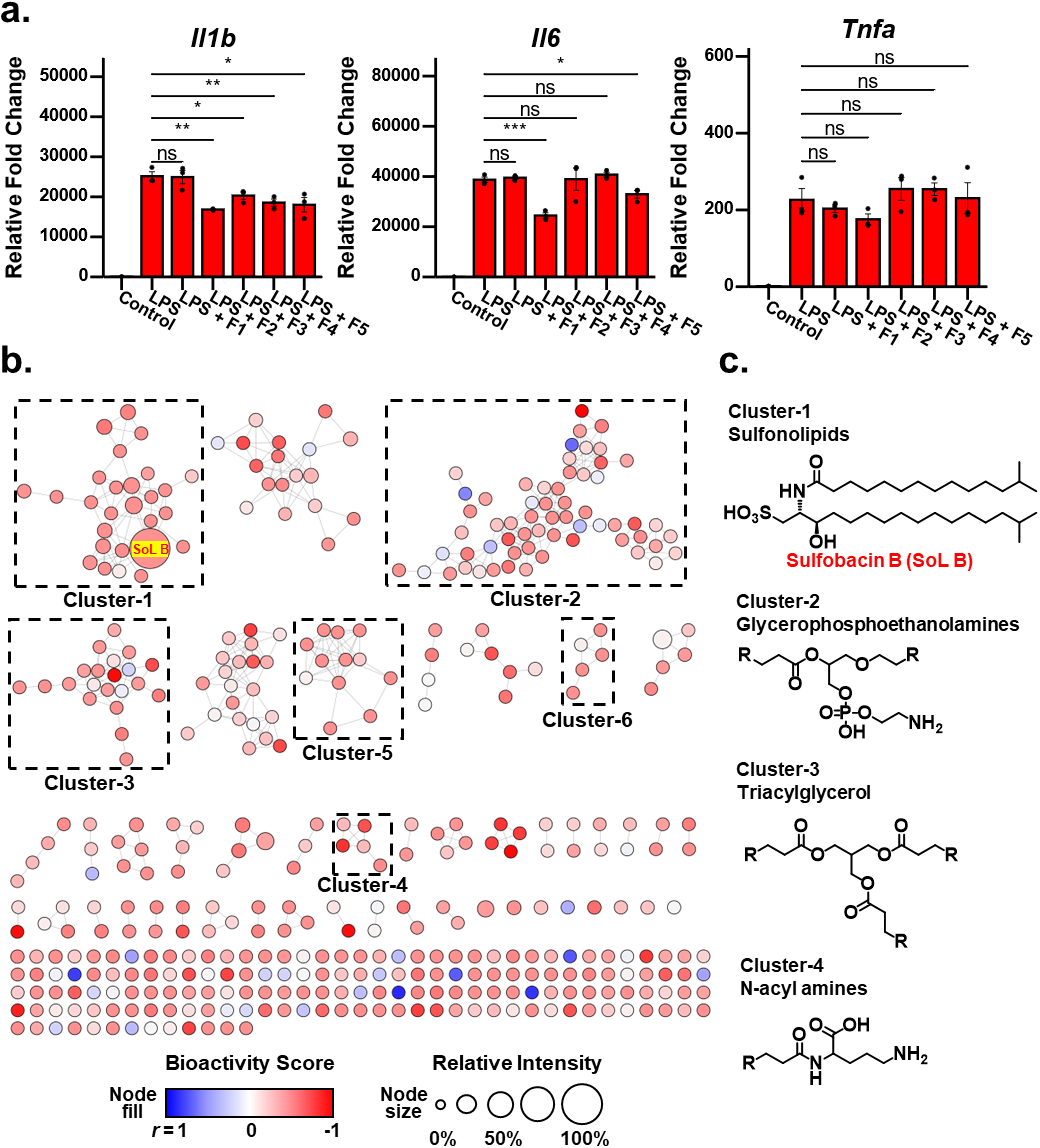
Bioactive molecular networking of *A. timonensis* OMV fractions suggests multiple compound classes contributing to an overall anti-inflammatory effect against LPS-induced inflammation. (a) Relative fold change expression pro-inflammatory cytokines *Il1b, Il6,* and *Tnfa* after treating mouse peritoneal macrophages 1 μg/mL LPS, or 1 μg/mL LPS with fractions of *A. timonensis* OMVs at 20 μg/mL (*n* = 3 wells per treatment). Fraction 2 (F2) showed the strongest suppression among other fractions and was further investigated for its bioactive components. Significance was determined using two-sided Student’s *t*-test. For all *p*-values, * 0.05 > *p* > 0.01, *** 0.001 > *p* > 0.0001, and ns: not significant. (b) Untargeted metabolomics and subsequent bioactive molecular networking analysis of Fraction 2 of *A. timonensis* OMVs revealed multiple clusters of molecules including SoLs which were abundant in this active fraction. (c) Predicted chemical structures of the major clusters, suggesting other potential contributors to the biological activity of Fraction 2.

To investigate these potential biologically active components, the molecular composition of F2 was analyzed by untargeted metabolomics using high resolution mass spectrometry (HRMS/MS) followed by molecular networking using the widely used Global Natural Product Social (GNPS) molecular networking tool.^50^ A bioactivity-based molecular network was then constructed by correlating the abundance change of molecular features with the expression change of inflammatory marker *Il1b* using spearman correlation.^51^ The correlative bioactivity score was mapped to the node color parameter while the abundance was mapped to the node size. Individual nodes represent molecular features in F2 and edges between nodes indicate similarity in the MS/MS fragmentation pattern of the nodes (**Figure 7b**). Interrogating the network, we identified several clusters with both promising bioactivity score and high chemoinformatic confidence scores, supporting the presence of bioactive components in F2. The cluster containing the largest node in F2 corresponded to SoLs, and the largest node corresponded to SoL B, which was confirmed by MS/MS comparison to our in-house standard (**Supplemental Figure S4**). Notably, in our previous study, pure SoL B was found to suppress LPS-induced inflammation by binding to and displacing LPS from the canonical binding pocket of the TLR4/MD-2 complex, supporting the observed anti-inflammatory activity of F2.^35^

To identify other potential contributors from F2, we further analyzed the HRMS/MS data using the chemoinformatic platform SIRIUS, which afforded computationally predicted chemical formulas based on exact mass measurements as well as predicted compound classes and putative structural identities based on MS/MS fragmentation patterns.^52–55^ In addition to confirming the known biologically active SoL cluster (**Supplemental Figure S5**), we prioritized five other clusters, clusters 2 through 6, representing families of potentially bioactive compounds and predicted their compound classes, including glycerophosphoethanolamines, triacylglycerols, *N*-acyl amines, glycerophosphoserines, and another family of N-acyl amines, respectively. Based on these classes, we further identified assigned structural features for clusters 2 through 4 (**Figure 7b,c**), then selected representative nodes for putative structural identifications based on their MS/MS fingerprint matching to known and predicted database spectra (**Supplemental Tables S2-S5**) (**Supplemental Figures S6-S8**).^53–55^ This result demonstrates the diverse chemical composition, especially lipid molecules, of *A. timonensis* OMVs. Furthermore, our observation that SoL B was a major compound in the bioactive molecular network with good bioactivity score suggests that other compounds with similar bioactivity scores may also contribute to the anti-colitic effect of *A. timonensis* colonization.

### Sulfonolipids isolated from *A. timonensis* OMVs delayed colitis progression in *Il10*-deficient mice

Our bioinformatic analysis suggesting SoLs, particularly SoL B, as a major contributing class to the observed anti-inflammatory activity aligns with the existing literature reporting that SoL B is an immunomodulatory lipid with anti-inflammatory effects on cells *in vitro* and in some *in vivo* applications.^35,56^ Thus, we were encouraged to determine if SoL B delivered by *A. timonensis*-derived OMVs plays a role in delaying colitis in the IL10 KO model. From the bioactive *A. timonensis* OMV F2, we further purified SoL B (**Supplemental Fig. S9a**) and determined this fraction to be free of LPS contamination using an LAL endotoxin assay (**Supplemental Fig. S9b**). We used this purified SoL B to treat IL10 KO mice (SoL B; n = 6) at a dosage of 1 mg/kg by intraperitoneal (I.P.) injection. As a vehicle control, IL10 KO mice (Veh.; n = 5) were treated with 1X PBS alongside the SoL B group. Concurrently with the first I.P. injection, both groups were switched to a piroxicam-containing diet to induce colitis. Mouse weight was measured over the course of 17 days, which revealed that mice in the SoL B group showed lower weight change at necropsy compared to mice in the Vehicle group (**Supplemental Fig. S10a**) and showed significantly lower inflammation in gross pathological parameters with a similarly trending but non-significant pattern in histological parameters (**Figure 8a**) (**Supplemental File 5**). In the distal colon, SoL B treatment further significantly suppressed the expression of pro-inflammatory markers *Tnfa* and *Lcn2*, while other cytokines *Il6* and *Nos2* showed lowered but non-significant outcomes (**Figure 8b-e**). In addition, treatment showed a reduced but non-significant expression of *Ifng*, *Il17a*, and *Il12b*, suggesting SoL B attenuated Th1 and Th17 adaptive immune responses in the host (**Supplemental Fig. S10b**). Taken together, these data indicate that SoL B treatment by I.P. injection at a dosage of 1 mg/kg can suppress colitis development in IL10 KO mice, supporting that SoL B likely plays a role in the anti-inflammatory phenotype observed with *A. timonensis* colonization.

**Figure 8.**
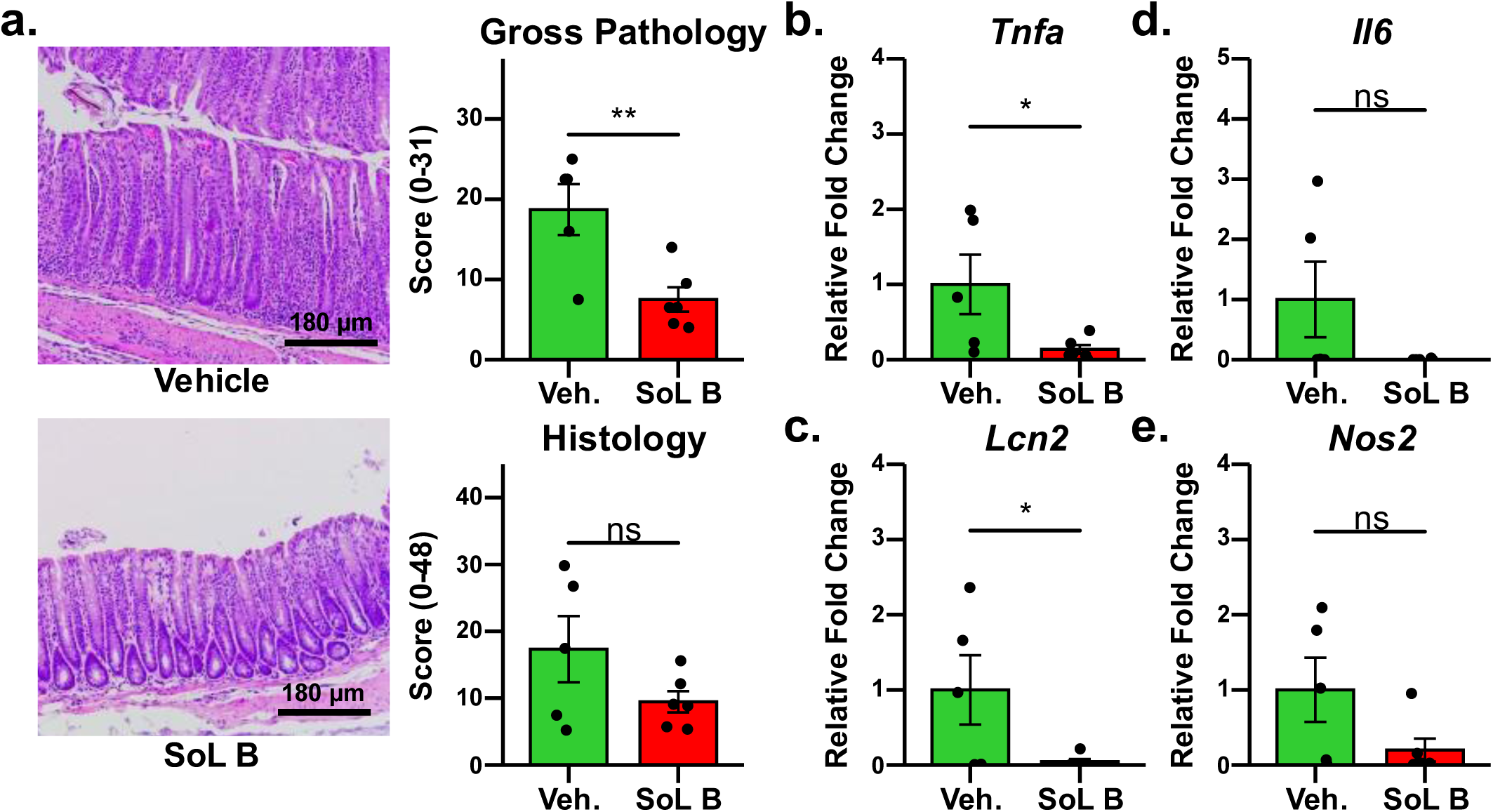
IL10 KO mice treated with purified SoL B by I.P. injection exhibit delayed colitis progression. (a) Representative histology images illustrate the difference in colon pathology associated with purified SoL B treatment (SoL B) compared to vehicle control (Vehicle). Gross pathological analysis revealed significantly lower levels of inflammation in SoL B-treated mice. Histological analysis also supports lower inflammation in the SoL B-treated mice. (b-e) RT-qPCR showed expression of pro-inflammatory markers were decreased in SoL B-treated mice (SoL B) compared to vehicle-treated mice (Veh.), supporting that SoL B treatment suppresses colitis development.

## Discussion

In this work, we demonstrate that colonization of *A. timonensis* DSM 27924 in an *Il10*-deficient (IL10 KO) mouse model of IBD suppresses colitis development which is related to host lipidome modulation rather than through counteracting microbial dysbiosis. We found that OMVs derived from the plasma of *A. timonensis*-colonized mice carry immunomodulatory SoLs leading to significantly increased SoL abundance in the plasma of *A. timonensis*-colonized mice. We then show that fractions of *A. timonensis* OMVs suppress LPS-induced inflammation and find immunomodulatory SoLs in the active fraction. We further leverage analytical tools integrating bioactivity and metabolomics to probe the chemical composition of the active *A. timonensis* OMV fraction, potentially expanding the biologically active metabolites of *A. timonensis* OMVs. Overall, we demonstrate that *A. timonensis* colonization in IL10 KO mice suppresses colitis progression via OMV-mediated delivery of immunomodulatory *A. timonensis* metabolites including SoLs to mouse plasma, potentially resulting in host lipidomic modulation.

*Alistipes* is a genus with emerging clinical relevance and there are conflicting reports about the impact of various *Alistipes* species on different diseases, particularly those relating to inflammation such as IBD.^16^ Several studies have negatively correlated *Alistipes* abundance in the gut microbiome to IBD, all suggesting that *Alistipes* may play a role in protecting against IBD.^20,35^ As such, we sought to explore the molecular mechanisms by which *Alistipes* affects the host in IBD. We used IL10 KO mice as a model of IBD since these mice are genetically predisposed to developing intestinal inflammation and disease onset in these mice can be synchronized by administration of the non-steroidal anti-inflammatory drug (NSAID) piroxicam.^33,34^ Using this colitis model, we observed that colonization with *A. timonensis* DSM 27924 by oral gavage resulted in suppressed colitis development through significantly reduced gross pathology and histology scores, as well as significantly reduced inflammatory cytokine expression and favorable modulation of tight junction protein expression. These results indicate that *A. timonesis* colonization attenuates colitis induced by chronic NSAID feeding.

Typically, this type of interaction is accompanied by major pathological restructuring of the gut microbiome, which can further exacerbate the host pro-inflammatory immune response.^57,58^ However, our microbial composition analysis by 16S rRNA sequencing revealed that *A. timonensis* colonization had little effect on the overall gut microbiome composition except for an increase in the abundance of *A. timonensis* itself. This result was unusual and intriguing, indicating that instead of inducing a compensatory microbial shift, *A. timonensis* alone directly influences the host to exert an anti-colitic effect, possibly through the modulation of host signaling pathways. Similarly, the human gut commensal *Bacteroides thetaiotaomicron* is known to produce a lipid molecule, α-galactosylceramide, that induces a host anti-cancer immune response through natural killer T cell activation.^59^

To investigate the influence of *Alistipes* on host immune signaling molecules, targeted lipidomic analysis of host circulating lipids revealed a significantly different lipid profile upon *A. timonensis* colonization characterized by increased phosphatidic acids and decreased triglycerides. The majority of differential lipids belonged to the phosphatidic acid and triglyceride classes. Notably, in acutely stimulated immune cells, arachidonic acid (AA) is mobilized first from membrane phospholipids, including phosphatidic acids, which is then replenished by triacylglycerol AA after inflammation has subsided.^45^ We observed a consistent trend in that AA-containing phosphatidic acids were accumulated in *A. timonensis*-colonized compared to uncolonized samples while triglycerides were decreased in *A. timonensis*-colonized samples. This may represent the replenishment of AA from triglycerides to phospholipids in an *A. timonensis*-mediated suppression of the inflammatory response in response to piroxicam challenge. Taken together, the clear shift in host lipidome profile alongside the changes in balance between AA-containing phospholipids and triglycerides support that *A. timonensis*-colonization indeed affects host immune signaling.

Since *A. timonensis* colonization did not induce a microbiome shift but did affect the profile of host circulating lipids, we considered if *A. timonensis* was producing and translocating immunomodulatory factors to the host circulation. Like other Gram-negative bacteria that produce OMVs as a mechanism to interact with their environments, *A. timonensis* also produces OMVs. OMVs are known to cross the mucosal barrier and epithelial cell layer, effectively delivering microbial factors to the host circulation.^28,31,32^ We found that OMVs isolated and purified from the blood of *A. timonensis*-colonized mice were enriched in SoL B, the major sulfonolipid of *A. timonensis* and a component of its outer membranes, suggesting that the purified OMVs were derived from *A. timonensis* and indicating that *A. timonensis* indeed delivers factors to the host circulation via OMVs.^47^ We noted that the fold-change increase of SoL B in the whole plasma of *A. timonensis*-colonized compared to uncolonized samples was significantly larger than that of the feces, indicating that SoL B is a non-gut restricted microbial metabolite and suggesting that *A. timonensis*-derived SoL B may be actively translocated from the gut to the host circulation by OMVs. Further HRMS/MS analysis of the whole plasma also revealed the increased abundance of SoLs A and F in *A. timonensis*-colonized blood samples, indicating that *A. timonensis* OMVs deliver other factors in addition to SoL B to the host circulation. Importantly, SoL A and B were previously identified to exert anti-inflammatory activity against LPS-induced inflammation, supporting that *A. timonensis* may mediate its anti-colitic effect through delivery of functional metabolites to the host circulation via OMVs.^35^

To determine if *A. timonensis* OMVs could recapitulate the anti-colitic phenotype of *A. timonensis* colonization, we fractionated *A. timonensis* OMVs into different fractions by normal phase chromatography, tested their biological activity, and pinpointed the component responsible for the activity. In an *in vitro* assay measuring LPS-induced pro-inflammatory cytokine expression, we found that Fraction 2 significantly suppressed the expression of *Il1b* and *Il6*. Because the inflammatory response in IL10 KO mice is characterized by leakage of gut contents, such as LPS, into the circulation, we hypothesized that Fraction 2’s ability to suppress LPS-induced inflammation in macrophages represents a mechanism by which *A. timonensis* OMVs suppress colitis after *A. timonensis* colonization. To identify bioactive components in the active Fraction 2, we conducted untargeted metabolomics and bioactive molecular networking, which revealed an abundance of SoL B that was negatively correlated with the expression of *Il1b*, supporting its hypothesized involvement in the observed anti-colitic phenotype. While our previous work supports the role of SoL B in the observed phenotype, our understanding of the complexity and intricacy of host-microbial interactions suggested that additional *A. timonensis*-derived components may also play a role.^35^ Thus, from the bioactivity-based molecular network, we prioritized five additional clusters with good bioactivity scores and numerous members for chemoinformatic analysis to predict compound classes and potential structure identities using SIRIUS, a machine learning-based chemoinformatics platform.^52–55^ Our analysis revealed several other lipid-type compound classes and matched spectral features to several compounds, many of which had similar or better bioactivity score than SoL B. This result significantly expands the pool of potentially bioactive metabolites delivered to host blood by *A. timonensis* OMVs and opens the door for further exploration of *A. timonensis*’ effect on inflammation. The presence of potentially bioactive metabolites in *A. timonensis* OMVs suggests that the intact OMVs may also exhibit biological activity. While this activity could be relevant to the effect of *A. timonensis* colonization in IL10 KO mice, OMVs are highly complex in nature. Thus, we focused our study on the identification of bioactive components and fractions of OMVs in the present study, which serves as a foundational step toward exploring the role of intact OMVs in *A. timonensis* colonization in subsequent investigations.

Finally, to confirm the role of SoL B in the observed anti-inflammatory phenotype with *A. timonensis* colonization, we purified SoL B from the active *A. timonensis* OMV fraction and tested its efficacy in an independent cohort of IL10 KO mice. We found that purified SoL B does suppress colitis in the IL10 KO model compared to vehicle-treated control mice. Compared to *A. timonensis* colonization, mice treated with SoL B saw an almost equivalent level of reduction in gross pathological parameters. This report is the first to demonstrate the ability of SoL B to suppress colitis in IL10 KO mice and supports that SoL B is a major contributor in mediating the anti-inflammatory response observed with *A. timonensis* colonization. Further, the presence of SoL B in *A. timonensis* OMVs coupled with its effectiveness *in vivo* strongly supports our hypothesis that *A. timonensis* modulates the inflammatory response in IL10 KO mice through delivery of immunomodulatory lipids by OMVs.

By establishing OMVs as a delivery vehicle for immunomodulatory SoLs and other potentially bioactive lipids, we show that *A. timonensis* can translocate functional metabolites from the gut into the plasma and affect the balance of circulating host immune signaling molecules. This provides a potential explanation for *Alistipes’* influences on distant organs such as the liver and cardiovascular system.^17,19^ For example, circulating *Alistipes* OMVs in the blood may directly deliver active metabolites to the liver or cardiovascular tissue inducing a long-range protective effect against liver and cardiovascular inflammation. While this study is focused on *Alistipes*, we envision that investigating the OMVs of other biologically relevant human gut microbiota may serve as a vital strategy to reveal additional biologically active non-gut restricted microbial metabolites in host-microbe interactions.

## Materials and Methods

### Bacterial strain cultivation

*Alistipes timonensis* DSM 27924 was obtained from the German Collection of Microogranisms and Cell Cultures (DSMZ). For the production of OMVs, we used *Alistipes timonensis* DSM 27924. *A. timonensis* was cultivated under anaerobic conditions on solid brain heart infusion agar plates or in liquid reinforced clostridium media at 37°C.

### Animals

*Il10*-deficient (IL10 KO) mice on the C57BL/6 background were originally purchased from The Jackson Laboratory. All mice were bred and maintained in specific pathogen free conditions at the University of South Carolina and were maintained on a 12-hour light/dark cycle with unlimited access to water and food (Inotiv Teklad, 8604). All animal protocols were approved by the University of South Carolina Institutional Animal Care and Use Committee.

### *Il10*-deficient mouse model and treatments

At 7.5-9 weeks of age, female IL-10 KO mice were divided into one of the following experimental groups: 1) piroxicam treated (*n* = 10 mice distributed into 3 cages); 2) piroxicam treated and colonized with *A. timonensis* (*n* = 11 mice distributed into 3 cages). Female control IL10 KO mice (*n* = 17 mice distributed into 6 cages) at 7.5-9 weeks old were caged separately from piroxicam-treated IL10 KO mice. Colitis development was initiated as previously described.^35^ Briefly, on day 0, all mice were switched to a diet supplemented with 100 ppm of piroxicam (Cayman Chemical, 13368; Inotiv Teklad, TD.210442). From day 0, mice in the first group were gavaged with RCM (reinforced clostridium media) to serve as a vehicle control, while *A. timonensis*-colonized mice were gavaged with 7.48×10^7^ CFU/mL of *A. timonensis* in RCM. All mice were gavaged three times per week for 2.5 weeks until necropsy at 17 days post piroxicam initiation.

For purified SoL B treatment, female IL10 KO mice at 8.5-10 weeks of age were divided into two groups: vehicle control (*n* = 5 mice distributed into 3 cages – originally 3 cages of 6 females, but 1 mouse died due to reasons unrelated to the model) and SoL B treated (*n* = 6 mice distributed into 3 cages). SoL B was prepared at a concentration of 0.2 mg/mL in 1X PBS with <1% DMSO and 100 µL was delivered to mice at a dosage of 1 mg/kg by I.P. injection. Control mice were injected with 100 μL of 1X PBS vehicle. On day 0, all mice were switched to a piroxicam-supplemented diet as above prior to the first I.P. injection. All mice were injected three times per week for 2.5 weeks until necropsy at 17 days post piroxicam induction.

Mouse body weight was tracked as an indicator for piroxicam-accelerated colitis progression. At necropsy, intestinal tissues were collected for evaluating colitis severity, intestinal contents were collected for microbiome and metabolite analysis, and plasma was collected for lipidomics and OMV detection. Colitis severity including gross pathology analysis, histopathology analysis, and proinflammatory cytokine gene expression were performed as previously described.^35^ The relative abundance of mammalian mRNA transcripts was calculated using the ΔΔ*C_T_* method and normalized to *Eef2* levels. The oligonucleotides used for qRT-PCR were: *Eef2 forward:* TGTCAGTCATCGCCCATGTG, reverse: CATCCTTGCGAGTGTCAGTGA*; Tnfa* forward: AGCCAGGAGGGAGAACAGAAAC, reverse: CCAGTGAGTGAAAGGGACAGAACC; *Nos2* forward TTGGGTCTTGTTCACTCCACGG, reverse: CCTCTTTCAGGTCACTTTGGTAGG*; Il6* forward; GAAATGATGGATGCTACCAAACTG, reverse: CTCTCTGAAGGACTCTGGCTTTG; *Il1b* forward: CTCAATGGACAGAATATCAACCAAC, reverse: GGCTGTGCCGTCTTTCATTAC.

### Immunofluorescence light microscopy

Fresh frozen colon tissues were cross-sectioned at 10 μm and fixed with 100% Acetone for 10 mins at −20°C before rinsing with PBS. Sections were blocked with 5% bovine serum albumin (BSA) for 1 h at room temperature. Sections were incubated with anti-claudin-3 or anti-claudin-7 antibody diluted at 1:100 in BSA overnight at 4°C. Sections were rinsed with PBS, and the secondary antibody ALEXA FLUOR^™^ 488 diluted at 1:600 in BSA was added to the sections for 1 h in the dark. Then, sections were rinsed with PBS and mounted with FluoroGel + DAPI. Sections were examined and photographed using a Zeiss Axio Imager M2 fluorescent light microscope equipped with Zen imaging software.

### Genomic DNA extraction and 16S rRNA gene sequencing

Genomic DNA (gDNA) was extracted from Control, uncolonized, and *A. timonensis*-colonized mouse feces using a QIAamp Fast DNA Stool Mini Kit (Qiagen). gDNA was quantified using a BioSpectrometer (Eppendorf) and sent to the University of Alabama Birmingham Microbiome Institutional Research Core for 16S rRNA sequencing. Sequencing data was processed and analyzed using Qiime2 (v2024.5) and R (v4.4.1) along with the R package Qiime2R to import Qiime2 objects to R.^60^

### Plasma lipidomic analysis

Flash frozen mouse plasma samples were sent to MetwareBio for lipid-focused extraction and targeted lipidomic analysis and quantification of host lipids. Fragmentation and retention time data corresponding to SoLs was provided to facilitate targeted analysis and quantification of detectable SoLs. Abundance data was analyzed using R (v4.4.1) along with the R package MetaboAnalystR (v4.0).^61^

### OMV purification

*A. timonensis* was cultivated as described above and OMVs were obtained following established isolation procedures with minimal modification.^62^ Briefly, bacterial cells were harvested by centrifugation at 12,000 x *g* for 30 minutes at 4°C. The supernatants were then filtered through 0.2 μm bottle top aPES membrane filters (Thermo Scientific) to remove residual bacterial cells. Filtered supernatants were then aliquoted into polycarbonate tubes for ultracentrifugation at 100,000 x *g* for 2 hours at 4°C. After ultracentrifugation, the supernatant was decanted, replaced with fresh filtered supernatant, and centrifuged again at 100,000 x *g* for 2 hours at 4°C until all the filtered supernatant was processed. The resulting OMV pellets were then resuspended in filter sterilized 1X phosphate buffered saline (PBS) pH 8.0 and passed again through 0.2 μm syringe filters to ensure bacterial cells had been completely removed. Diluted OMV samples were then pooled and concentrated in Spin-X UF 20 100 kDa PES membrane centrifugal filters (Corning). Protein concentrations of the purified OMV samples were determined using a Bradford assay and samples were prepared for electron microscopy at 200 ug/mL protein concentration.

### Density gradient purification of OMVs from mouse plasma

667 μL aliquots of plasma from *A. timonensis*-colonized IL10 KO mice were added to 3.333 mL of 60% iodixanol (OptiPrep) stock to create a 50% iodixanol OMV load solution. 4 mL of 40%, 30%, 20%, and 10% Iodixanol solutions were prepared in 1X PBS. Iodixanol solutions were then layered in order of decreasing Iodixanol concentration, beginning with the 50% iodixanol OMV load solution, into polycarbonate tubes for ultracentrifugation, and 1 mL 1X PBS was added to the top of each sample. The samples were then ultracentrifuged at 100,000 x *g* for 18 hours at 4°C, forming a continuous density gradient. After ultracentrifugation, each sample was fractionated into 21 1 mL fractions, taken from the top. The presence of SoLs in each fraction was determined by HPLC-MS and HPLC-MS/MS analysis. SoL-containing fractions were then pooled and concentrated in Spin-X UF 20 100 kDa PES membrane centrifugal filters (Corning).

### Size Exclusion Chromatography of OMVs from mouse plasma

Concentrated OMV solutions were then briefly centrifuged to remove cell debris and aggregates, before being subjected to Size Exclusion Chromatography. Apex 4B Size Exclusion Columns (Everest Biolabs) were washed with 10 mL of 1X PBS before 0.5 mL of concentrated OMVs were applied to each column. The first 2 mL of eluate was discarded, and the following eight 0.5 mL fractions were collected. OMV-containing fractions were determined via Bradford Assay and confirmed by HPLC-MS/MS targeting SoL B. OMV-containing fractions were then pooled and stored at - 20°C prior to analysis.

### TEM visualization

10 μL aliquots of OMV samples either undiluted or prepared at 200 μg/mL protein concentration were treated with 10 μL of 4% OsO4 (ThermoFisher Scientific) for five minutes. Then 10 μL of the treated OMV sample was loaded onto copper formvar/carbon coated 200 mesh grids and allowed to settle for 5 minutes. Excess solution was dabbed away and the grids were negative stained with 10 μL of 5% uranyl acetate (ThermoFisher Scientific) for 5 minutes before excess solution was dabbed away. Grids were allowed to dry and samples were viewed using a Hitachi HT7800 TEM (Hitachi High-Tech Co.).

### Dynamic light scattering of OMV samples

1mL aliquots of OMV samples from plasma or *A. timonensis* culture were prepared at 100 μg/mL protein in 1X PBS and added to plastic cuvettes. Particle size distribution was then measured by dynamic light scattering using a Zetasizer Nano ZS (Malvern Panalytical). Experiments were performed in triplicate and all measurements were repeated three times.

### OMV fractionation and SoL isolation

Approximately 200 μg of OMVs were dried and lyophilized prior to extraction with methanol. The methanol extract was deposited on silica and dried overnight prior to loading for open normal phase column chromatography. The OMV extract sample was eluted using five column volumes each of 15:1, 7:1, 5:1, 3:1, and 1:1 dichloromethane:methanol which was collected and dried by rotary evaporation and by lyophilization. Fraction samples were prepared at 20 ug/mL in dimethylsulfoxide for subsequent *in vitro* assays and HRMS/MS analysis. SoL B was purified from OMV fractions and tested for LPS contamination as previously described.^35^

### Preparation of primary mouse macrophages

Primary mouse macrophages were prepared by first introducing 3 mL of 3% thioglycolate to mice via intraperitoneal injection. After 3 days, 10 mL of chilled PBS was introduced intraperitoneally to flush out macrophages. The cell suspension was then separated by centrifugation at 300 x g for 5 min. Cells were seeded in culture dishes containing DMEM with 10% FBS for 1 hour before being rinsed with serum-free DMEM two times to remove unattached cells.

### Cell treatment

Samples of *A. timonensis* OMV fractions or *A. timonensis* OMV fractions + LPS were added to the culture media of primary mouse macrophages for 6 hours before the treatment media was removed. The cells were then washed twice with 1X PBS and detached from the cell culture plate with 0.25% trypsin. Cells were removed from the suspension by centrifugation and split equally for mRNA extraction and metabolite extraction. *A. timonensis* OMV fractions were prepared at 20 ug/mL and LPS was prepared at 1 ug/mL.

### mRNA extraction and RT-qPCR

Cells were lysed with TRI Reagent (Zymo Research) and total RNA was extracted from the cell lysate using a Direct-Zol^TM^ RNA MiniPrep Kit (Zymo Research) according to the manufacturer’s protocol. The quality and quantity of RNA was then determined using a Nanodrop One (ThermoFisher) and 1,000 ng of mRNA from each sample was reverse-transcribed to cDNA using iScript cDNA Synthesis Kit (Bio-Rad). qPCR was conducted on a CFX96 system (Bio-Rad) using iQ SYBR Green Supermix (Bio-Rad). All primers used for qPCR analysis were synthesized by Integrated DNA Technologies. The 18S RNA was used as a housekeeping gene. The relative amount of target mRNA was determined using the comparative threshold (Ct) method by normalizing target mRNA Ct values to those of 18S RNA. PCR thermal cycling conditions were 3 min at 95°C, and 40 cycles of 15 s at 95°C and 58 s at 60°C. Samples were run in triplicate. Relative gene expression levels were calculated using the ΔΔ*C_T_* method and expression levels of 18S were used to normalize the results.

### HPLC-MS/MS analysis of OMV fractions and mouse plasma

OMV samples were extracted by addition of an equal volume of methanol. The organic solvent was then removed and dried under a gentle stream of N_2_. The residue was reconstituted in methanol for HPLC-MS/MS analysis using a Vanquish HPLC (ThermoFisher Scientific) coupled to a Q-Exactive HF-X hybrid Quadrupole-Orbitrap mass spectrometer using electrospray ionization in positive mode. Analytical scale HPLC separation was performed on a Waters Xbridge BEH column (2.1 x 100 mm) using a basic mobile phase consisting of solvent A: 60:40 acetonitrile:water + 0.1% formic acid + 10 mM ammonium formate and solvent B: 10:90 acetonitrile:isopropanol + 0.1% formic acid + 10 mM ammonium formate delivered at a flow rate of 0.2 mL/min in a gradient starting from 37% B to 98% B over 20 minutes, followed by 98% B for 5 minutes to wash the column, back to 37% B over 3.5 minutes, and finally 37% B for 7.5 minutes to re-equilibrate the column. MS scans were obtained in the orbitrap analyzer which was scanned from 200 to 2000 *m/z* at a resolution of 60,000. MS/MS was conducted using data-dependent acquisition with a resolution of 30,000, isolation window of 2.0 *m/z*, and dynamic exclusion time of 15 seconds.

### Bioactive Molecular Networking and chemoinformatic analysis

HPLC-HRMS/MS data was processed using MZmine3 following the GNPS FBMN workflow with minimal changes.^63^ Molecular networks were constructed using the quickstart GNPS FBMN setting with no changes.^50^ Raw data used for this analysis was deposited in the University of California, San Diego Center for Computational Mass Spectrometry MassIVE database (https://doi.org/doi:10.25345/C55M62K1G). Bioactivity scores were assigned using a custom R script which calculated Pearson correlation coefficients between each molecular feature and the activity of each fraction.^51^ Finally, bioactive molecular networks were visualized in Cytoscape v3.9.1.^64^ Representative nodes were selected for further chemoinformatic analysis with SIRIUS v6.0.6 using the CANOPUS and CSI:FingerID tools to predict compound classes as well as representative node structures and substructure annotations.^52,53,55^

### Statistics and reproducibility

For all experiments, sample sizes were chosen to achieve statistical significance, and data from outliers were excluded based on Grubbs’ test for outliers. For animal experiments, randomization was applied to assign animals to groups and cages and blinding was applied in the analysis of histopathology data.

## Author contributions

E.A.O., M.K.M., M.E., and J.L. designed the study. E.A.O., M.K.M., A.C., X.L., M.M., Y.W., L.E.V., T.Z., B.V., and R.T. conducted the study and acquired the data. E.A.O., M.K.M., M.E., and J.L. analyzed the data. E.A.O. and J.L. drafted the manuscript. All authors interpreted the data, read, and critically revised the manuscript, and approved the final version.

## Acknowledgements

We acknowledge Drs. Michael Walla and William Cotham of the University of South Carolina Mass Spectrometry Facility for their assistance with HRMS/MS data acquisition and processing. We also acknowledge Dr. Diego Altomare, the director of the University of South Carolina Microarray Core Facility for his help with high throughput RT-qPCR for measuring cytokine expression, as well as Mr. Cole Espinosa for his assistance in *A. timonensis* OMV purification. We acknowledge the Instrumentation Resource Facility at the University of South Carolina School of Medicine for their assistance with processing and staining intestinal tissues for histopathology analysis. We acknowledge the University of Alabama Birmingham Microbiome Institutional Research Core for their assistance with 16S sequencing of IL10 KO mouse fecal samples.

## Disclosure statement

No potential conflict of interest was reported by the author(s).

## Data availability statement

All data generated or analyzed during this study are included in the published article and its supplementary information files. Raw MS/MS data used for the bioactivity-based molecular networking analysis was deposited in the University of California, San Diego Center for Computational Mass Spectrometry MassIVE database (https://doi.org/doi:10.25345/C55M62K1G). Raw MS/MS data was also published alongside raw 16S sequencing data in Zenodo (https://doi.org/10.5281/zenodo.13987894).

## Funding

This work was partially supported by the National Institutes of Health (NIH) through Grant 1R35GM150565 awarded to J.L. and another NIH Grant 1R01AI184916 awarded to M.E.

## Notes

### Competing Interest Statement

The authors have declared no competing interest.

### Summary of Updates

This version of the manuscript has been revised based on Reviewers' comments.

